# Convolutional neural networks for mild diabetic retinopathy detection: an experimental study

**DOI:** 10.1101/763136

**Authors:** Rubina Sarki, Sandra Michalska, Khandakar Ahmed, Hua Wang, Yanchun Zhang

**Affiliations:** Institute for Sustainable Industries & Liveable Cities, Victoria University, Melbourne, Victoria, Australia

## Abstract

Currently, Diabetes and the associated Diabetic Retinopathy (DR) instances are increasing at an alarming rate. Numerous previous research has focused on automated DR detection from fundus photography. The classification of *severe* cases of pathological indications in the eye has achieved over 90% accuracy. Still, the *mild* cases are challenging to detect due to CNN inability to identify the subtle features, discrimnative of disease. The data used (i.e. annotated fundus photographies) was obtained from 2 publicly available sources – Messidor and Kaggle. The experiments were conducted with 13 Convolutional Neural Networks architectures, pre-trained on large-scale ImageNet database using the concept of Transfer Learning. Several performance improvement techniques were applied, such as: (i) fine-tuning, (ii) data augmentation, and (iii) volume increase. The results were measured against the standard Accuracy metric on the testing dataset. After the extensive experimentation, the maximum Accuracy of 86% on No DR/Mild DR classification task was obtained for *ResNet50* model with fine-tuning (un-freeze and re-train the layers from 100 onwards), and RMSProp Optimiser trained on the combined Messidor + Kaggle (aug) datasets. Despite promising results, Deep learning continues to be an empirical approach that requires extensive experimentation in order to arrive at the most optimal solution. The comprehensive evaluation of numerous CNN architectures was conducted in order to facilitate an early DR detection. Furthermore, several performance improvement techniques were assessed to address the CNN limitation in subtle eye lesions identification. The model also included various levels of image quality (low/high resolution, under/over-exposure, out-of-focus etc.), in order to prove its robustness and ability to adapt to real-world conditions.

## Introduction

Approximately 420 mln people worldwide have been diagnosed with Diabetes [1], and its prevalence has doubled in the past 30 years [2]. The number of people affected is only expected to increase, particularly in Asia [3]. Nearly 30% of those suffering from Diabetes are expected to develop the Diabetic Retinopathy (DR) – a chronic eye disease that is considered a leading cause of vision loss among working-age adults [1, 4]. The eventual blindness resulting from DR is irreversible, though it can be prevented through regular fundus examinations [5].

Effective treatment is available for patients identified through *early* DR identification [6]. Needless to say, a timely detection of pathological indication in the eye leading to DR is critical. It not only allows to avoid the late invasive treatments and high medical expenses, but most importantly – to reduce the risk of potential sight loss. The manual methods of diagnosis prove limiting given the worldwide increase in prevalence of both Diabetes and its retinal complications [7]. Currently, the ophthalmologist-to-patient ratio is approx. 1:1000 in China [5]. Furthermore, the traditional approaches reliant on human assessment require high expertise, as well as promote inconsistency among the readers [1]. Labour and time-consuming nature of manual screening services has motivated the development of automated retinal lesions detection methods, in particular early stages of DR.

Deep Neural Network model is a sequence of mathematical operations applied to the input, such as pixel value in the image [8], where the training is performed by presenting the network with multiple examples, as opposed to unflexible rule-based programming underlying the conventional methodologies [9]. Deep learning, in particular Convolutional Neural Networks (CNN), has been widely explored in the field of DR detection [10–14],largely surpassing previous image recognition methodologies [14]. Overall, Deep learning has demonstrated tremendous potential in healthcare domain, enabling the identification of patients likely to develop a disease in the future [6]. In terms of DR, the applications range from binary classification (No DR/DR), to multi-level classification based on condition severity scale (No DR/Mild DR/Moderate DR/Severe DR). CNNs, with their multi-layer feature representations, have already shown outstanding results in discovering the intricate structures in high-dimensional datasets. The models have proven successful at learning the most discriminative, and often abstract aspects of the image, while remaining insensitive to irrelevant details such as orientation, illumination or background.

The numerous challenges in automatic DR detection have been identified in literature. The diagnosis is particularly difficult for patients with early stage of DR [1]. As highlighted by Pratt et al. [12], Neural Networks struggle to learn sufficiently deep features to detect intricate aspects of Mild DR. In the same study, approx. 93% of mild cases were incorrectly classified as healthy eye instances. The problem is illustrated on Fig 1, displaying various stages of DR advancement, and the associated features visibility. The numerous accuracy improvement techniques such as dimensionality reduction or feature augmention have been proposed in literature. Still, the studies on Deep learning-based DR detection consistently report high performance on severe cases, while identification of mild cases still remains a challenge. This limitation impede the wider application of fully automated mass-screening due to potential omission of early phase of DR, leading to more advanced condition development in the future. Also, according to the study conducted by Ting et al. [15], *the referable stage* of DR (mild) was 5x more prevalent than *the vision-threatening stage* of DR (severe), demonstrating the significance of early lesions detection.

**Fig 1.**
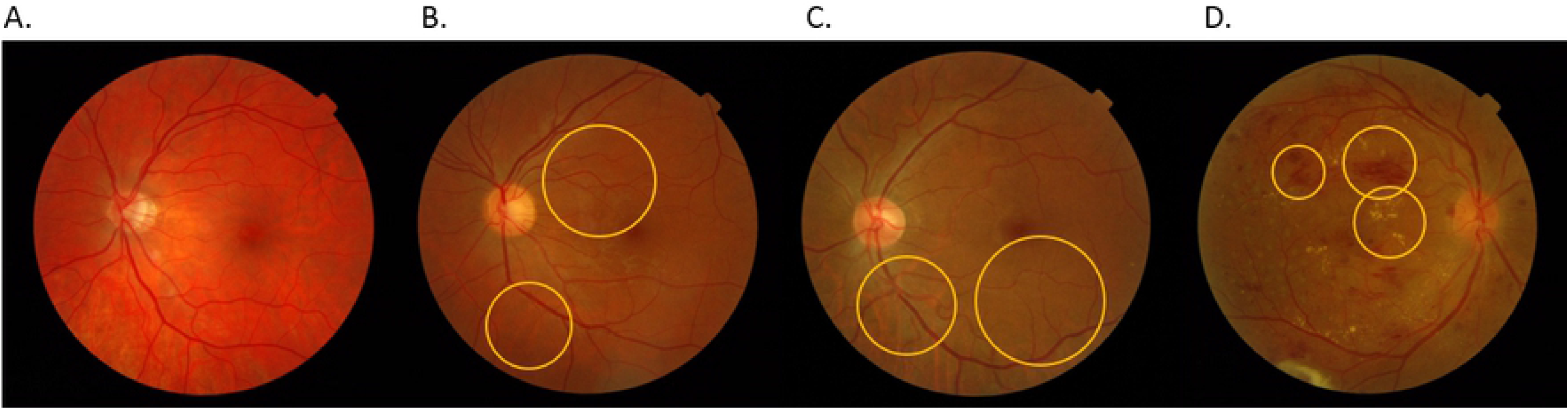
The examples of different stages of DR advancement on fundus images. (A) No DR – healthy retina; (B) Mild DR – abnormal growth of blood vessels and ‘cotton wool’ spots formation (early indication); (C) Moderate DR – abnormal growth of blood vessels and ‘cotton wool’ spots formation (mid-stage indication); (D) Severe DR – hard exudates, aneurysms and hemorrhages (advanced indication).

Transfer learning has already been validated and demonstrated promising results in medical image recognition. The concept uses knowledge learned on primary task, and its re-purpose to secondary task. Transfer learning is particularly useful in Deep learning applications that require vast amount of data and substantial computational resources. The state-of-the-art CNN models, pre-trained on the large public image repository have been used as part of this study, following the concept of Transfer learning. Using the weights initialised, the top layers of Neural Networks have been trained for customised No DR/Mild DR binary classification from publicly available fundus image corpora. The improved classification performance via Transfer learning has already been reported in prior research on automated DR detection [16]. Unlike previous approaches, the study conducted focuses entirely on Mild DR instances – currently challenging to identify. The several task-specific data augmentation techniques for classification performance improvement are further evaluated. Finally, the fully-automated Deep learning-based system facilitates methodology reproducibility and consistency in order to streamline an early DR detection process, increasing the access to mass-screening services among the population-at-risk.

## Materials and methods

### Study design

The overarching aim of the study is the performance improvement of early DR detection from fundus images of Mild DR and healthy retina through the extensive experimentation. The associated objectives can be identified as follows:

- Comprehensive evaluation of 13 CNN architectures using concept of Transfer learning;
- Models fine-tuning to reflect the specifics of the application case-study;
- Various Optimisers performance assessment and the optimal one selection;
- 2 datasets combination and augmentation for further accuracy improvement.

To illustrate the steps followed, the high-level process pipeline is presented in Fig 2.

**Fig 2.**
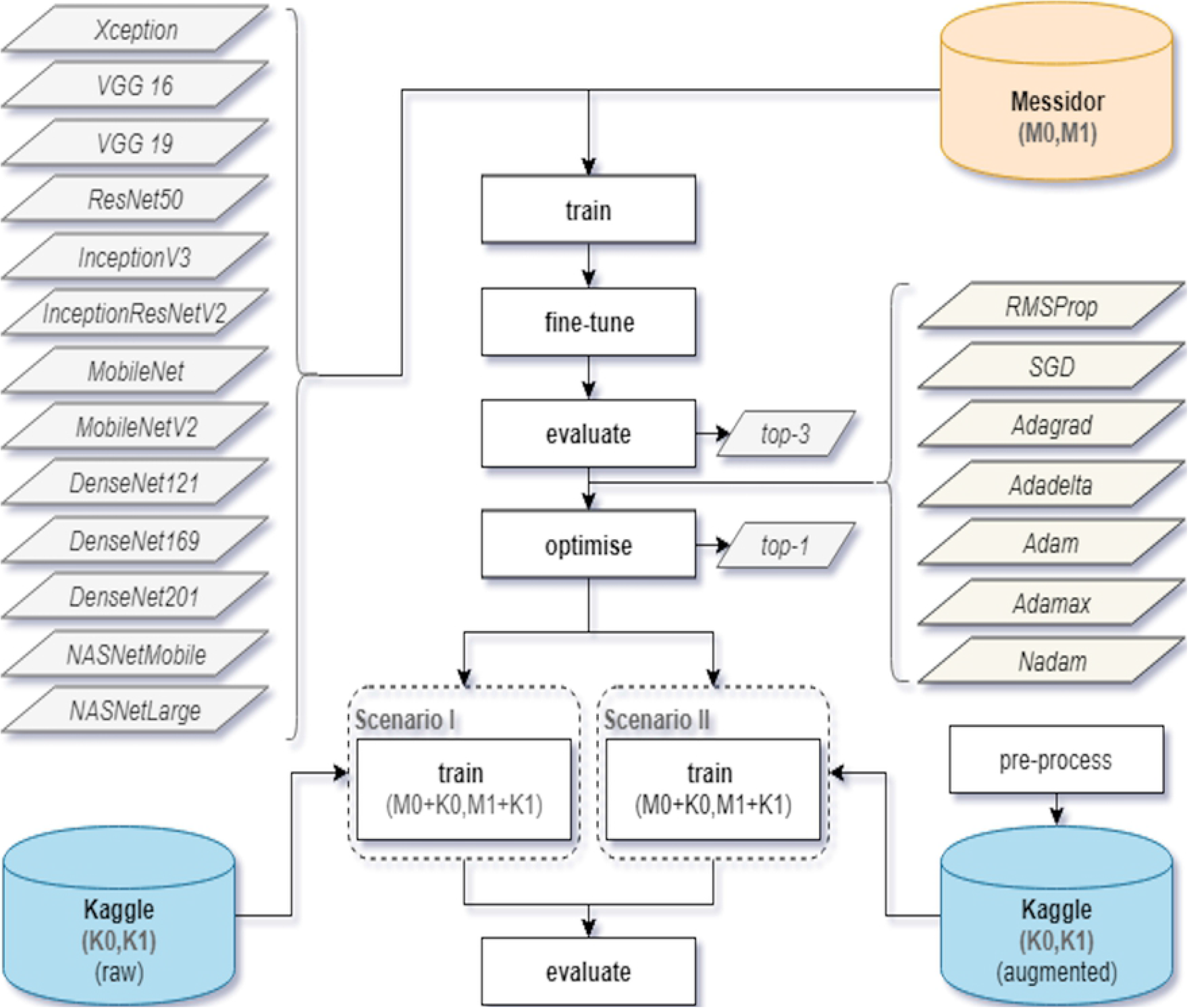
The high-level process pipeline.

### Data collection

The data was acquired from publicly available corpora, i.e. Kaggle and Messidor. Kaggle dataset contains 35126 fundus images, annotated for 5-class identification (No DR, Mild DR, Moderate DR, Severe DR, Proliferative DR), while Messidor dataset contains 1200 fundus images, annotated for 4-class identification. Both datasets include colour photographs of right and left eye. The images dimensions vary between low-hundreds to low-thousands. The quality of data differs significantly between the datasets. Messidor, despite its relatively small scale, is considered a high fidelity source with reliable labelling, while Kaggle consists of a large number of noisy and often misannotated images. The raw Kaggle data more closely reflects real-world scenario, where images are taken under different conditions, thus resulting in various quality levels. The challenge lies in the potential eye lesions detection despite the observed noisiness in application dataset. Fig 3 demonstrates the comparative data distribution for Messidor and Kaggle datasets among the respective DR classes (the exact numbers of images for each dataset/class can be found in Table 2).

**Fig 3.**
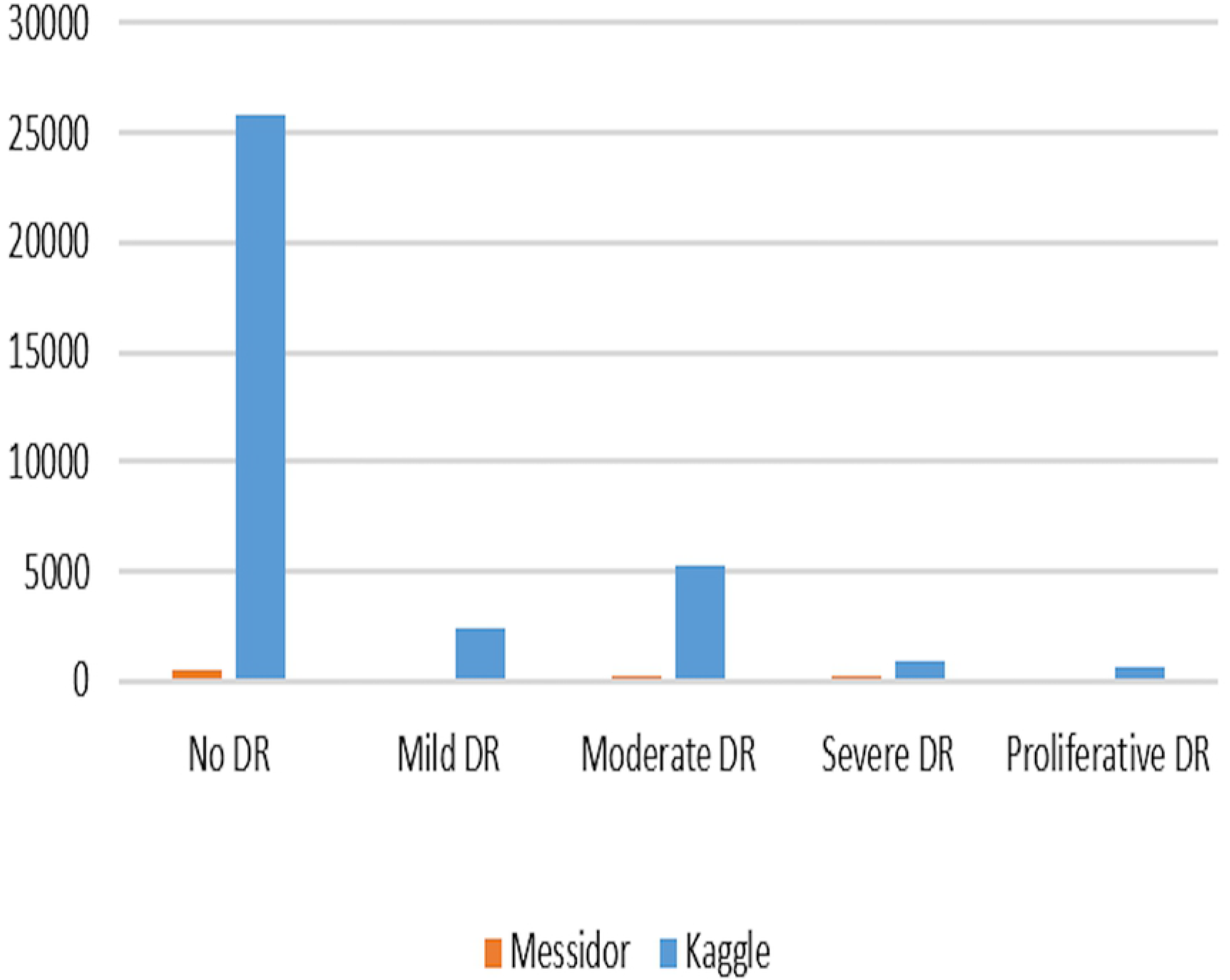
Data distribution for Messidor and Kaggle datasets.

### Transfer learning

The knowledge transfer from primary to secondary task frequently acts as an only solution in highly-specialised disciplines, where the availability of large-scale quality data proves challenging. The adoption of already pre-trained models is not only the efficient optimisation procedure, but also supports the classification improvement. The first layers of CNNs learn to recognise the generic features such as edges, patterns or textures, whereas the top layers focus on more abstract and task-specific aspects of the image, such as blood vessels or hemorrhages. Training only the top layers of target dataset, while using the initialised parameters for the remaining ones is commonly employed approach, in particular in computer vision domain. Apart from efficiency gains, fewer parameters to train also reduce the risk of overfitting, which is a major problem in Neural Networks training process [12]. The CNN models used in the experiments along with their characteristics are presented in Table 1.

**Table 1.**
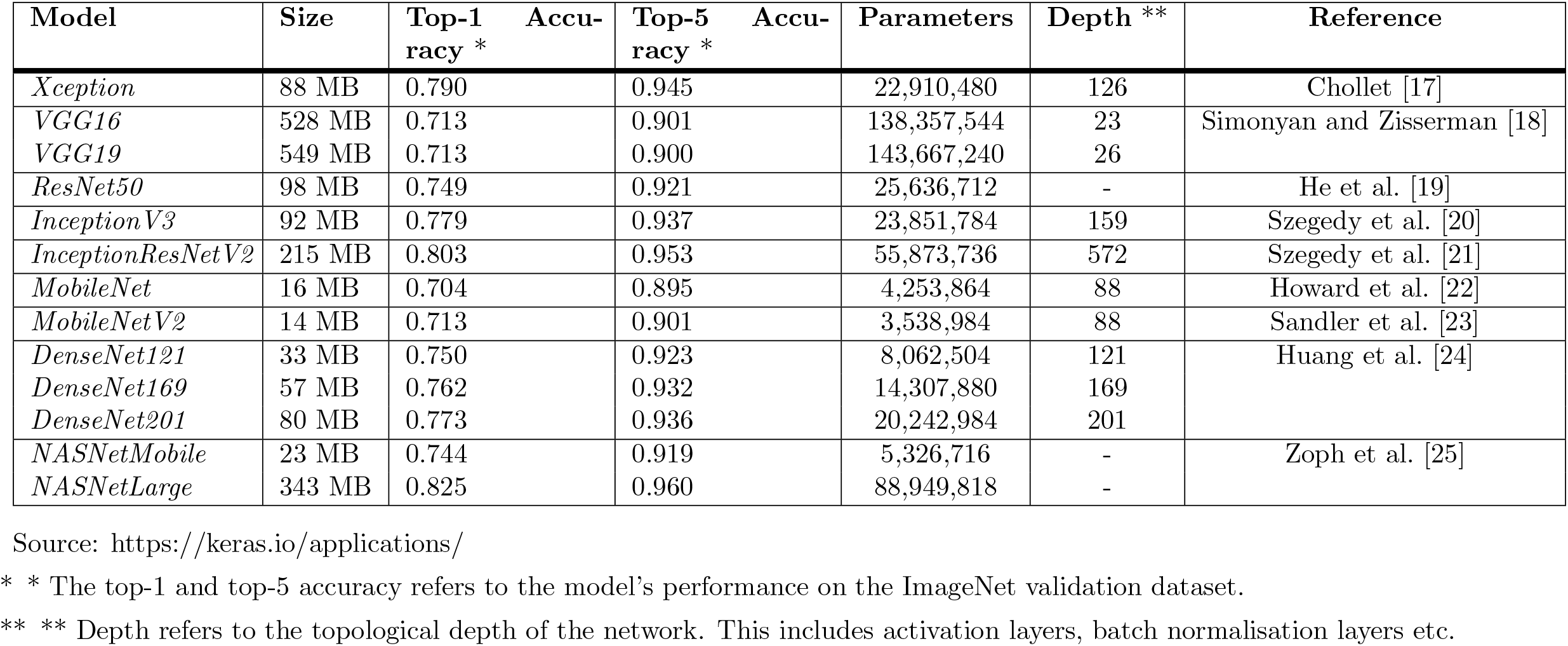
The CNN models pre-trained on ImageNet and their characteristics.

**Table 2.** The DR severity levels according to ETDRS and the number of images used in each experiment.

| Severity level | Id | Description | M | K | K | M + K | M + K |
| --- | --- | --- | --- | --- | --- | --- | --- |
|  |  |  |  | (raw) | (aug) | (raw) | (aug) |
| *No DR* | 0 | No abnormalities. | 546 | 25810 | 50976 | 26356 | 51522 |
| *Mild DR* | 1 | Microaneurysms only. | 153 | 2443 | 55410 | 2596 | 55563 |
| *Moderate DR* | 2 | More than just microaneurysms, but less than severe NPDR. | 247 | 5292 | - | - | - |
| *Severe DR* | 3 | Any of the following and no signs of proliferative retinopathy: (1) severe intraretinal hemorrhages and microaneurysms in each of four quadrants; (2) definite venous beading in two or more quadrants; (3) prominent IRMA in one or more quadrants. | 254 | 873 | - | - | - |
| *Proliferative DR* | 4 | One or both of the following: (1) neovascularisation; (2) vitreous/preretinal hemorrhage. | - | 708 | - | - | - |

Legend: M - Messidor, K - Kaggle, NPDR - Non-Proliferative Diabetic Retinopathy, IRMA - Intraretinal Microvascular Abnormalities.

### Experiment setting

The algorithms were implemented using Keras library^1^, with TensorFlow^2^ as a back-end. The images resolution has been standardised to a uniform size in accordance with input requirements of each model. The number of epochs, i.e. complete forward and backward passes through the network, was set to 20 due to the already pre-trained models use. The training/testing split was set to 60/40 given the small-to-moderate dataset size. The stratified random sampling was performed in order to ensure the correct class distribution and final findings reliability. The mini-batch size was set to 32, and the cross-entropy loss function was selected due to its suitability for binary classification. The default Optimiser was RMSProp. The standard evaluation metric of Accuracy on testing dataset was used for final results validation.

### Performance improvement

#### Fine-tuning

The CNN models adopted in the study were pre-trained on a large-scale ImageNet dataset that spans numerous categories includings flowers, fruits, animals etc. Such models obtain high performance on classification tasks for the objects present in the training dataset, while prove limiting in their application to niche domains, such as DR detection. Diagnosis of pathological indications in fundoscopy depends on a wide range of complex features, and their localisations within the image [1]. In each layer of CNN, there is a new representation of input image by progressive extraction of the most distinctive characteristics [10]. For example, the first layer is able to learn edges, while the last layer can recognise exudates – a DR classification feature [1]. As a result, the following scenarios were considered in the experiments: (1) <u>only</u> the top layer removal and network re-train (the existing approaches); and (2) the *n* top layers removal and network re-train (the proposed approach). The parameter *n* vary across the CNNs used, and depends on the total number of layers present in each model structure. The threshold of 100 was selected, and the subsequent layers of each model were ‘un-frozen’ and fine-tuned to the application dataset. The initial 100 layers were treated as a fixed feature extractor [26], while the remaining layers were adapted to specific characteristics of fundus photography. The potential classification improvement on DR detection task was evaluated as a result of the proposed models customisation. In the study conducted by Zhang et al. [5], the performance accuracy of Deep learning-based DR detection system improved from 95.68% to 97.15% as a result of fine-tuning.

#### Optimisation

During the training process, the weights of Neural Network nodes are adjusted accordingly in order to minimise the loss function. However, the magnitude and direction of weights adjustment is strongly dependent on the Optimiser used. The most important parameters that determine the Optimiser’s performance are: Learning rate and Regularisation. Too large/too small value of Learning rate results in either non-convergence of the loss function, or in the reach of the local, but not absolute minima, respectively. At the same time, the Regularisation allows to avoid model overfitting by penalising the dominating weights values for the correct predictions. Consequently, the classifier generalisation capability improves, when exposed to the new data. The Optimisers used in the experiments were as follows: *(1) RMSprop*, *(2) SGD*, *(3) Adagrad*, *(4) Adadelta*, *(5) Adam*, *(6) Adamax* and *(7) Nadam*.

#### Volume

Deep learning benefits from high-volume data. The larger number of both No DR as well as Mild DR instances significantly increases model’s reliability and allows for more distinctive patterns detection. Thus, the previously-used small-scale Messidor dataset has been combined with large-scale Kaggle dataset, i.e. the respective No DR and Mild DR classes have been merged together. Table 2 presents the numbers of images used in each scenario along with the descriptions of the particular DR stages. The severity scale used is in accordance with the Early Treatment Diabetic Retinopathy Study (ETDRS) [5]. The horizontal line separates No DR (M0,K0) and Mild DR (M1,K1) from all cases available in Messidor and Kaggle datasets.

#### Augmentation

Compared to the Messidor dataset, the Kaggle dataset consists of larger proportion of low fidelity data. The images were captured with different fundus cameras, resulting in various quality levels. The relatively noisy character of images is observed through their blurriness, under/over-exposure, presence of unrelated artifacts, and so on. The raw format of Kaggle dataset closely reflects the nature of DR detection in real-world settings, where substantial variability in data quality is observed between the institutions.

In order to evaluate the potential classification improvement due to pre-processing techniques applied, the following steps have been performed: *(1) crop*, *(2) resize*, *(3) rotate* and *(4) mirror*. The example of the original image, and the augmentation steps implemented are presented in Fig 4. *Cropping and resizing* (1 + 2) allows to focus on pathological indications with greater level of detail, which proves important for DR discrimination. Additionally, the subsequent *rotating and mirroring* (3 + 4) substantially expands the dataset, alleviating the imbalance issues between the classes. The comparison of images volumes before and after augmentation is included in Fig 5.

**Fig 4.**
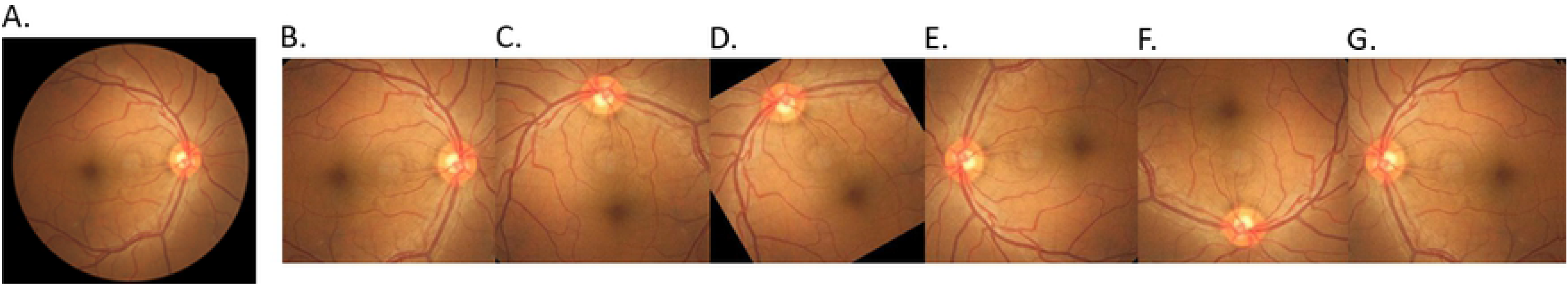
The examples of data augmentation steps performed on Mild DR fundus image from Kaggle dataset. (A) Original; (B) Crop; (C) Rotate 90°;(D) Rotate 120°; (E) Rotate 180°; (F) Rotate 270°; (G) Mirror.

**Fig 5.**
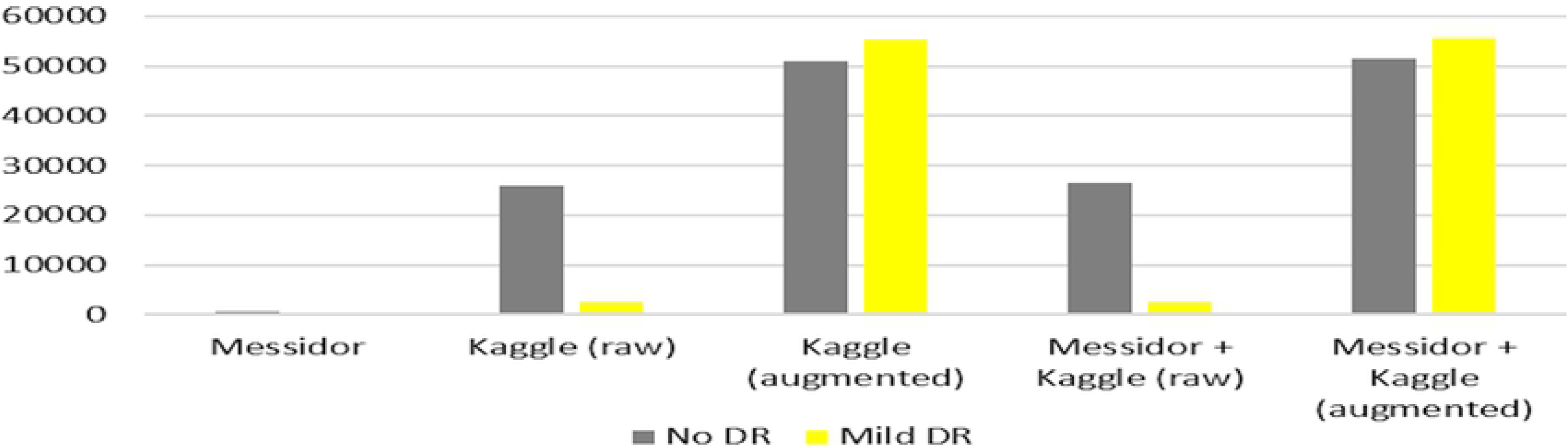
Data distribution before and after augmentation for No DR and Mild DR classes.

## Results

### Models comparison

The 13 pre-trained CNNs were compared in terms of yielded Accuracy on testing dataset (Table 3). Additionally, the fine-tuning was applied as an alternative to the default option. After removal and re-training of *n* layers from 100 onwards (*n* was CNN-dependent), the performance obtained for each model was used for comparison purposes. The fine-tuning effect was calculated in terms of percentage Accuracy increase/decrease. Then, the maximum Accuracy was selected for each model (either default or after fine-tuning). Finally, the top 3 CNN architectures with highest classification performance on Messidor dataset progressed to the subsequent optimisation procedure (Fig 2).

The Accuracy after each epoch was further plotted in order to investigate the models convergence capabilities in the default and fine-tuning scenario (Fig 6). As a result, the computational intensity was additionally evaluated.

**Table 3.** The accuracy comparison of pre-trained CNN models.

| Model | Accuracy | Accuracy (F-T*) | F-T* effect | Accuracy (max) |
| --- | --- | --- | --- | --- |
| Xception | 0.809 | 0.809 | ±00.0% | 0.809 |
| VGG16 | 0.809 | 0.809 | ±00.0% | 0.809 |
| **VGG19** | **0.813** | **0.813** | ±00.0% | **0.813** |
| **ResNet50** | **0.813** | **0.816** | +00.4% | **0.816** |
| InceptionV3 | 0.806 | 0.795 | −01.3% | 0.806 |
| InceptionResNetV2 | 0.812 | 0.582 | −28.4% | 0.812 |
| MobileNet | 0.583 | 0.556 | −04.7% | 0.583 |
| MobileNetV2 | 0.781 | 0.656 | −16.0% | 0.781 |
| DenseNet121 | 0.795 | 0.778 | −02.2% | 0.795 |
| DenseNet169 | 0.569 | 0.642 | +12.8% | 0.642 |
| DenseNet201 | 0.797 | 0.795 | −00.2% | 0.797 |
| NASNetMobile | 0.799 | 0.802 | +00.4% | 0.802 |
| **NASNetLarge** | **0.813** | **0.809** | −00.4% | **0.813** |

\* Fine-Tuning

**Fig 6.**
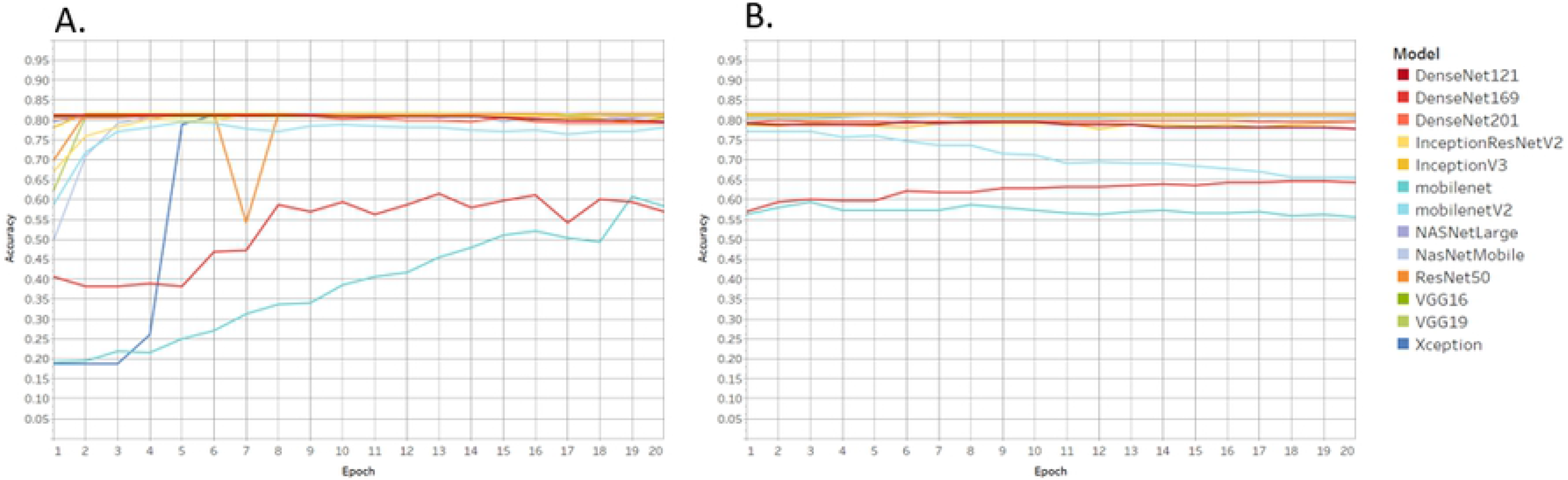
The validation accuracy achieved for the respective epochs. (A) Default; (B) Fine-tuning.

### Performance improvement

Following the top 3 CNN models selection (Table 3), the 7 most common in Deep learning applications Optimisers were evaluated as part of the hyper-parameters optimisation procedure. The most robust Optimiser in terms of validation accuracy for each of the 3 models was indicated. The highest performing model+Optimiser was selected for further augmentation process. The respective classes of both Messidor and Kaggle datasets (M0+K0,M1+K1) were merged together and used to train the max-accuracy model, as determined. The increase in dataset volume was expected to contribute towards performance improvement. Next, the augmentation of imbalanced low quality Kaggle data was conducted to further evaluate impact of image pre-processing on classification accuracy. The results of both scenarios (i.e. (I) Messidor + Kaggle (raw) data; and (II) Messidor + Kaggle (augmented)) are presented in Table 5.

## Discussion

### Automated DR detection

Increasing life expectancy, popular indulgent lifestyles and other contributing factors indicate that the number of people with Diabetes is projected to raise [12, 27]. This in turn places an enormous amount of pressure on available resources and infrastructure [28]. For instance, most of the patients with DR in China often neglect their condition, and fail to secure timely interventions resulting in severe state development [5]. Early identification of pathological indications effectively prevents further condition aggravation, and its impact on the affected individuals, their families, and associated medical expenses. Thus, the DR detection system allows to either (i) fully-automate the eye-screening process; or (ii) semi-automate the eye-screening process. First option requires sufficient level of accuracy, equivalent to that of retinal experts. According to British Diabetic Association guidelines, a minimum standard of 80% sensitivity and 95% specificity must be obtained for sight-threatening DR detection by any method [29]. Second option allows to downsize the large-scale mass-screening outputs to the potential DR cases, followed by human examination. Both scenarios significantly reduce the burden on skilled ophthalmologists and specialised facilities, making the process accessible to wider population, especially in low-resource settings.

### Performance improvement

The first part of the experiment included feature extraction initialised via Transfer learning using the pre-trained CNN models, followed by the removal of the top layer (existing approach). The comprehensive evaluation of total of 13 CNN architectures (including state-of-the-art) was performed. In the second part, the N layers were ‘un-frozen’ (over the threshold of 100), and subsequently re-trained to better adapt to the specifics of the application case-study (proposed approach). The combination of Messidor and Kaggle datasets was performed to further support model generalisation, given the variety of images provided for system training, as well as to benefit the model performance due to higher volume of training examples. The size of data used in training greatly affects the outcome of Neural Networks process [9]. The numerous pre-processing steps were implemented to measure potential accuracy improvement for No DR/Mild DR image classification. As Mild DR proves extremely challenging to identify from healthy retina due to only subtle indications of the disease, the data augmentation undertaken was believed to enhance pathological features visibility (e.g. zoom and crop).

The top 3 CNN architectures with the top layer removed and re-trained were *VGG19*, *ResNet50* and *NASNetLarge*, yielding the accuracy of 81.3% (Table 3). The lowest performance was obtained by *DenseNet169* (56.9%) and *MobileNet* (58.3%), respectively. In terms of *DenseNet169*, the characteristic feature of its structure that connects each layer to every other layer in a feed-forward manner did not prove to enhance the performance on No DR/Mild DR classification task. As for *MobileNet*, the results only confirmed its intended purpose for mobile applications due to its lightweight and streamlined architecture, which comes at a cost of the Accuracy.

The effect of fine-tuning (un-freezing the layers from 100 onwards) differed across the models. The Accuracy improvement observed was only minor, suggesting the relative suitability of pre-trained models to DR detection task. In other words, the CNN models were able to identify Mild DR from healthy retina despite being trained on un-related pictures from ImageNet repository. When no Accuracy increase is achieved, the further layers un-freezing is not recommended due to unnecessary computational time and cost incurred.

To complete the analysis on the effect of fine-tuning, the graphs depicting each CNN architecture performance at the respective epochs (single pass of the full training set) has been performed, as illustrated in Fig6. Despite no major influence on the classification Accuracy, faster model’s convergence was observed due to the fine-tuning applied. The higher number of layers un-frozen and re-trained made the models more task-specific, leading to an improved use of resources due to reduced training time for the most optimal performance. The finding was particularly noticable for the following models *Xception*, *MobileNet* and *DenseNet169*.

Next, the various Optimisers have been evaluated on the top 3 CNN architectures. Table 4 presents the Accuracy outputs obtained for each model. While there was no major impact on the classification performance for *VGG19*, the higher variability was observed for *ResNet50*, proving its sensitivity to the most suitable Optimiser selection. Overall, RMSProp proved the most optimal choice for 2 out of 3 models.ResNet50+RMSProp was selected as the max-Accuracy model+Optimiser option for No DR/Mild DR classification task.

**Table 4.**
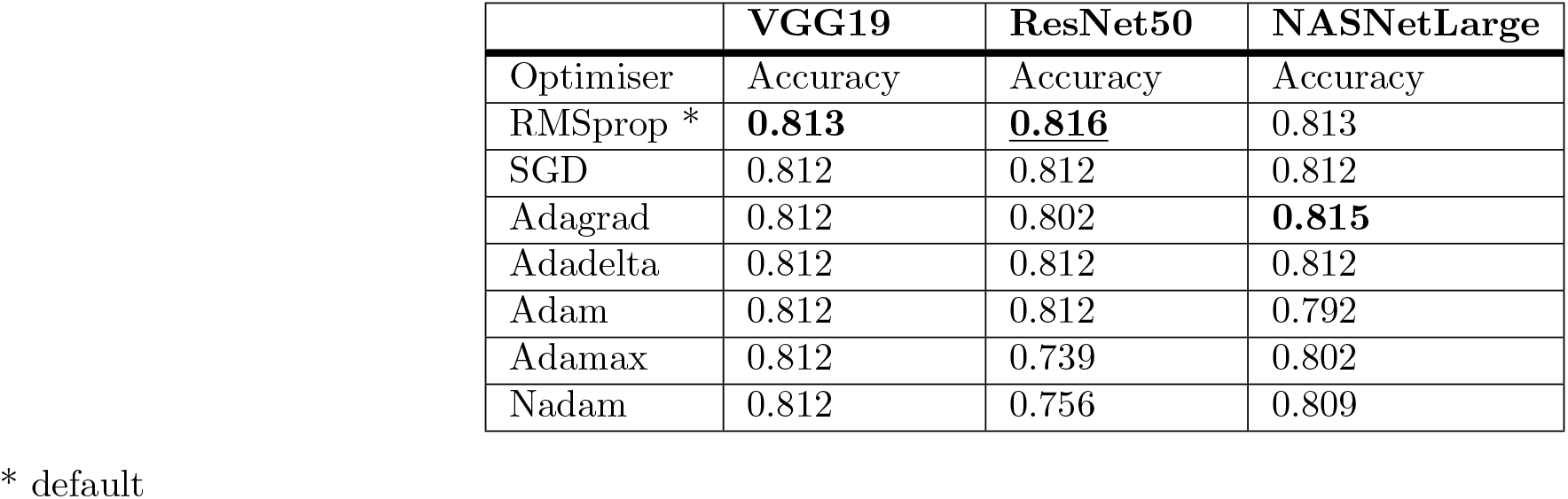
The Optimisers performance evaluation.

**Table 5.**
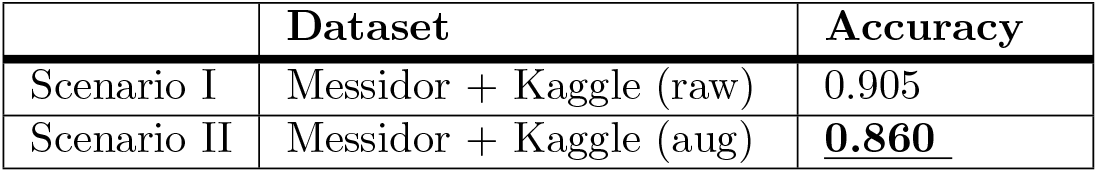
The effect of volume increase and data augmentation.

|  | Dataset | Accuracy |
| --- | --- | --- |
| Scenario I | Messidor + Kaggle (raw) | 0.905 |
| Scenario II | Messidor + Kaggle (aug) | **0.860** |

As the last step, the 2 scenarios were considered, namely (i) volume increase and (ii) data augmentation. As expected, the datasets combination, i.e. Messidor + Kaggle (aug) further improved the classification Accuracy by +5% (86%). Where as, combination of the datasets, i.e. Messidor + Kaggle (raw) results (90.5%) which is consider as case of data imbalance. Therefore, the augmented images helps in data imbalance, and did introduce greater variability of training examples required for the increased model performance and the improved generalisability. As a result, the max classification Accuracy on No DR/Mild DR classification task was achieved for *ResNet50* model with fine-tuning and RMSProp Optimiser trained on the combined Messidor + Kaggle (aug) datasets.

### Approach limitations

The several shortcomings of the study have been identified. Firstly, due to limited availability of Mild DR images, only small-to-moderate dataset size was used in the study. As a compensation procedure, the CNN models pre-trained on large-scale ImageNet database have been adopted. The appropriate data augmentation techniques have been further applied to expand the dataset, i.e. rotation, horizontal/vertical flip, etc. Secondly, the default hyper-parameters were followed, while training the classifiers (i.e. batch size, dropout, loss function, etc.). These are considered the best practice within the field. Still, the experiments with various Optimisers have been performed. Finally, the ‘black-box’ nature of Deep learning-based solution is frequently criticised, causing resistance in wider approach adoption by the practitioners. Models learn the features given the input data, and associate them with labels provided, the exact factors affecting the final prediction are not transparent and straightforward. According to Wong et al. [6], the major mindset shift is required in how clinicians and patients entrust clinical care to machines. The methodology also highlights the importance of accurate annotation process, as directly impacting the classifier performance. While Messidor dataset has been verified and labelled by trained ophthalmologists, the numerous mis-annotated images can be found in Kaggle dataset.

### Future work

As it is the initial study focusing on binary No DR/Mild DR classification, future work will cover finer-grained information extraction from cases previously identified as Mild DR. For instance, upon sufficient data availabilty, the model will allow to recognise the particular lesions such as exudates or aneurysms. The more in-depth classification will further assist the retinal practitioners in more efficient eye-screening procedure. Also, the highly varied input data (e.g. in terms of ethnicity, age group, level of lighting) will support the model robustness and flexibility. Additionaly, different scenarios with respect to the number of layers and nodes will be experimented with using TensorBoard for dynamic visualisation. Increased convolution layers are perceived to allow the model to learn deeper features [9, 12]. This in turn will enable the most optimal CNN architecture design (depth and width of the network) for maximum classification accuracy. An increase network dimensionality is the most direct way to enhance model performance [16]. Future work will also place more emphasis on outputs visualisation in order to obtain greater insight into the models internal workings, and further improve the classification capability. In particular, the identification of exact image regions that are associated with specific classification results will be highlighted, as well as the magnitude of each feature intensity (so called attention/saliency maps [8]). Improved understanding of the algorithm workings will facilitate the automated system wider adoption and acceptance among physicians [6]. Finally, the experiments with ensembling approach will be conducted, where the results of Neural Networks models trained on the same data will be averaged in order to evaluate further classification accuracy gains.

## Conclusion

Early detection and immediate treatment of DR is considered critical for irreversible vision loss prevention. The automated DR recognition has been a subject of many studies in the past, with main focus on binary No DR/DR classification [12]. According to the results, an identification of moderate to severe indications do not pose major difficulties due to pathological features high visibility. The issue arises with Mild DR instances recognition, where only minute lesions prove indicative of the condition, frequently undetected by the classifiers. Mild DR cases prediction is further challenged by the low quality of fundus photography that additionally complicates the recognition of subtle lesions in the eye. Thus, the study proposed the system that focuses entirely on Mild DR detection among the No DR instances, as unaddressed sufficiently in prior literature. Given the empirical nature of Deep learning, the numerous performance improvement techniques have been applied (i.e. (i) fine-tuning, (ii) data augmentation, and (iii) volume increase). Additional benefit of Deep learning incorporates the automatic features detection that are most discriminative between the classes. Such approach allows to avoid the shortcomings associated with empirical, and often subjective manual feature extraction methods. Furthermore, the study used the combined datasets from various sources to evaluate system robustness in its ability to adapt to the real-world scenarios. As stated by Wan et al. [16], the single data collection environment poses difficulty in accurate model validation. The system successfully facilitates the streamlining of labour-intensive eye-screening procedure, and serves as an auxiliary diagnostic reference whilst avoiding human subjectivity.

## Footnotes

1 https://keras.io/

2 https://www.tensorflow.org/guide/keras

